# Rapid adaptation to high temperatures in *Chironomus riparius*

**DOI:** 10.1101/205195

**Authors:** Quentin Foucault, Andreas Wieser, Ann-Marie Waldvogel, Barbara Feldmeyer, Markus Pfenninger

## Abstract

Effects of seasonal or daily temperature variation on fitness and physiology of ectothermic organisms and their ways to cope with such variations have been widely studied. However, the way multivoltines organisms cope with temperature variations from a generation to another is still not well understood and complex to identify. The aim of this study is to investigate whether the multivoltine midge *Chironomus riparius* Meigen (1803) responds mainly via acclimation as predicted by current theories, or if rapid genetic adaptation is involved. To investigate this issue, a common garden approach has been applied. A mix of larvae from five European populations was raised in the laboratory at three different pre-exposure temperatures (PET): 14, 20, 26°C. After three and five generations respectively, larvae were exposed to three treatment temperatures (TT) 14, 20, 26°C, mortality was monitored for the first 48h and after emergence. After three generations significant mortality rate differences depended on an interaction of PET and TT. This finding supports the hypothesis that chironomids respond rapidly to climatic variation via adaptive mechanisms, and to a lesser extent via phenotypic plasticity. The result of the experiment indicates that three generations were sufficient to adapt to warm temperature, decreasing the mortality rate, highlighting the potential for chironomids to rapidly respond to seasonally changing conditions.

## Introduction

Ambient temperature variation is a major factor affecting the fitness of organisms by regulating the speed of metabolic processes, and thus, everything from development to reproduction (Atkinson, 1994). Thus local thermal regimes impose strong selection pressures on organisms (Merilä & Hendry, 2014; Waldvogel *et al*., 2018). Ectothermic organisms are particularly affected, due to their dependence of body temperature from ambient temperature (Clarke & Fraser, 2004; Deutsch *et al*., 2008). In ectotherms, metabolic rates show a more or less linear temperature response over a large temperature range, however below and above a species specific threshold physiological performance rapidly drops. Similarly, the fitness of ectotherms displays an exponential increase with temperature. The mortality curve takes on a U shape, skewed towards higher temperatures with a maximal fitness at a certain, optimal temperature (T_opt_). The edges of this curve are characterised by a steep drop of survival at a certain low and high temperature (T_crit_) that usually coincides with the drop in physiological performance. Therefore, temperature related mortality in ectotherms seems mostly driven by physiological limitations (Pörtner, 2001; Paaijmans *et al*., 2013; Verberk *et al*., 2017). In temperate areas, seasonal and daily temperature variation selects for organisms with broad thermal tolerances following Janzen’s climate variability hypothesis (Janzen, 1967; Shah *et al*., 2017). This can be achieved e.g. by morphological modifications of body size depending on temperature to adjust energy requirements, known as Bergmann’s rule (French *et al*., 1998, Gardner *et al*., 2011), via both physiological plasticity and genetic adaptation (Atkinson & Sibly, 1997; Angilletta Jr *et al*., 2004). An example for a purely physiological organismic response to short exposure periods of both cold or heat stress are hardening mechanisms involving protein modifications at the cell level (Bowler, 2005; Overgaard *et al*., 2005). The effect of temperature variation on life history traits such as growth (Hauer & Benke, 1991; Frouz *et al*., 2002), reproduction (Péry & Garric, 2006) and metabolic processes (Edwards, 1958; Sankarperumal & Pandian, 1991; Hirthe *et al*., 2001) are well known. Even though the effects of short term temperature variations on physiology and fitness have been well studied, the processes governing responses to short term seasonal temperature variations on ectothermic organisms is still not fully understood (Paaijmans *et al*., 2013). This is in particular true for short lived, multivoltine species whose successive generations throughout the year are exposed to highly different temperature regimes, at least in temperate regions with their pronounced seasonality.

To cope with such a broad temperature variation, likely exceeding their optimal temperature range at least in the extremes, multivoltine ectothermic organisms can respond in different ways, such as (i) behavioural changes, allowing organisms to escape or mitigate the environmental pressure (Hutchison & Maness, 1979; Lencioni, 2004), (ii) phenotypic plasticity, resulting in changes in gene expression to alleviate abiotic stress (Johnston & Wilson, 2006) and/or (iii) by genetic adaptation (Bergland *et al*., 2014) to maximise their fitness. It is commonly assumed that i) and ii) act rather on short time scales, while iii) was presumed to proceed over long time-scales (Carroll *et al*., 2007). However, it becomes increasingly clear that evolutionary adaptation can be very rapid (Messer & Petrov, 2013), and can even occur on seasonal time-scales (Bergland *et al*., 2014).

Because both phenotypic plasticity and rapid adaptation are necessarily associated with changes in the phenotype, distinguishing whether phenotypic change is genetically based or results from phenotypic plasticity is difficult. It requires experimental testing over several generations in order to identify which of those mechanisms are involved (Merilä & Hendry, 2014). If the response to different temperatures is governed by phenotypic plasticity, we expect the phenotype and the resulting fitness of each generation to depend only on the temperature experienced by the according generation, independent of the temperature experienced by previous generations. Thus, irrespective of generation, individuals should perform similarly well in each temperature (i.e. according to the respective reaction norm for this temperature). If however, genetic adaption occurred, the phenotypic fitness response to different temperatures will rather depend on the temperature experienced by the previous generation(s), i.e. organisms should perform better in temperatures their progenitors were exposed to. Consequently, we expect an interaction between temperature experienced by previous generations and temperatures experienced by the test individuals.

The aim of this study is to infer how an ectothermic multivoltine invertebrate, *Chironomus riparius* (MEIGEN 1803)(Fig. 1), spending most of its life in an aquatic larval stage (Armitage *et al*., 1995), copes with short term temperature variations over a few generations - via phenotypic plasticity or rapid adaptation. The species fares very well with experimental culture conditions over several generations (Downe & Caspary, 1973; Nowak *et al*., 2007a; OECD, 2004; 2010) and is therefore well suited for evolutionary experiments. Moreover, the adaptive potential of this species to a large gradient of local climate conditions has already been shown (Nemec *et al*., 2013; Waldvogel *et al*., 2018). Wild populations of *C. riparius* harbours ample genetic variation and have a large effective population size (~10^6^, (Oppold & Pfenninger, 2017)). In addition, the several hundred offspring per breeding pair likely render selection processes on quantitative traits very effective (Pfenninger, 2017). In this study, we hypothesize therefore that *C. riparius* can genetically adapt rapidly within a few generations to different temperature regimes, which would give first experimental evidence for rapid adaptation to seasonal temperature fluctuations.

**Fig 1:**
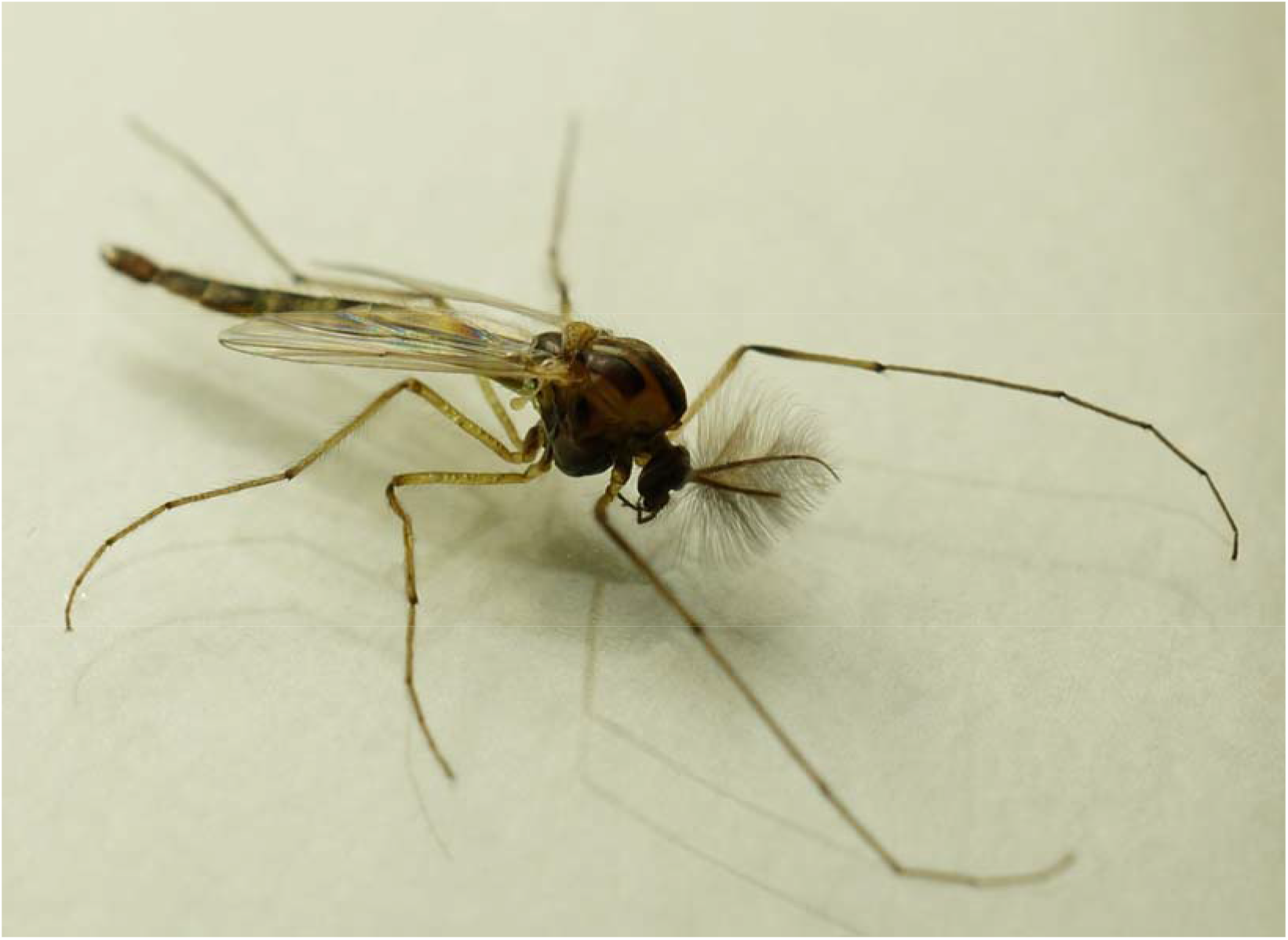
Macro-photography of an adult male *Chironomus riparius*

## Material and Methods

This experiment was performed on an admixed *C. riparius* culture composed of individuals from five different populations originating from Metz (NMF) and Lyon (MF) in France, Hasselroth in Germany (MG), Collobiano in Italy (SI) and Las Vegas in Spain (SS) (Oppold *et al*., 2016) (Supporting information 1). These populations were kept under laboratory conditions for a minimum of three generations prior to admixture. We decided to mix individuals from multiple populations in order to compensate for the expected loss of variation due to laboratory selection effect during the experiment (Nowak *et al*., 2007b), and to make sure that alleles involved in adaptation to the complete temperature gradient are present in the gene pool. From this admixed base population, which was kept for one generation at 20°C, three sub-populations were created. Two replicates from each subpopulation were raised at three different temperatures: 14 °C, 20 °C and 26 °C with light–dark rhythm of 16:8 h at 60% humidity, following the OECD guideline 219 (OECD, 2004; Oppold *et al*., 2016). These temperatures were oriented at the maximum, average and minimum mean temperature recorded during the meteorological summer (26.9°, 20.9° and 15.3°C) for the five populations of origin (Waldvogel *et al*., 2018). Those values were adjusted to obtain constant 6°C differences among experimental treatments. Field measured temperatures have been used to avoid reaching critical temperatures that are not commonly experienced in the wild, avoiding possible bias on the recorded mortality by artificially causing bottlenecks in the population (Supporting information 2).

Larvae were raised in medium constituted of purified water with sea salt (TropicMarin) adjusted to a conductivity of 520–540 μS cm^−1^ and pH 8. The bottom of the glass bowl (20 cm diameter) was covered with washed sand. Populations were raised at these temperatures for three and five generations to simulate the possible seasonal exposure range (i.e number of generations which could be expected during meteorological summer); this phase will hereafter be referred to as “Pre-Exposure Temperature” (PET).

### Survival (Mortality) test

Survival tests were common garden experiements performed in the third and fifth generation, i.e. individuals were exposed to PET for three and five generations, respectively. These two different time points were used to investigate possible acclimation or adaptation during the expected high and low number of generations possible during the meteorological summer. Generation time at the respective temperature was determined during the first generation, as time from the hatching of the eggs until the death of the adults. This was necessary, because generations started overlapping with the second generation, making it impossible to infer the exact beginning and end of a generation. The generation times used were inferred from their emergence time calculated in laboratory experiments: 33,6 days at 14°C, 18,1 days at 20 °C and 11,4 days at 26 °C (Oppold *et al*., 2016).

For each replicate, five egg clutches were put to hatch. Individuals coming from the same egg clutch will be referred to as families. 18 larvae from each family of each replicate were raised at three different experimental temperatures called “Treatment Temperature” (TT) for the analysis (18 larvae × 5 families × 3 temperatures × 2 population replicates) (Supporting information 3). Each larva was individually raised in six well plates (Ø3.5×2cm) filled with six mL of medium for 48 hours without feeding to avoid altering the medium. Larval mortality rates were measured first after 24 and then 48 hours. After 48 hours, the surviving larvae were pooled by families in glass bowls (Ø20 ×10 cm) with sediment and medium and reared until emergence. The mortality of pupation and emergence was calculated as the number of individuals not emerged per family, for each combination of PET and TT, when imagines were removed from bowls. During this stage, larvae were fed daily with dried fish food (0.5 mg/individual of grounded TetraMin® flakes) and the water level adjusted daily with deionized water in order to conserve the physico-chemical parameters.

### Statistics

Statistical analyses were performed using R (Version 3.2.3) in addition with Rstudio (Version 0.99.903). The normality of the data set was tested using qqplot, Kolmogorov-Smirnov tests and homoscedasticity as well with Levenes test. A linear mixed model was used to investigate the effect of the experimental factors on the mortality with the families as random factor followed by one-way ANOVA tests. In case of significant interactions of two or more factors, each instance of the interaction was analysed separately. In order to investigate significant differences between the TTs, ANOVA followed by Tukey *post-hoc* test were used for data following the assumptions of normality and homoscedasticity. Kruskal-Wallis tests followed by Dunn *post-hoc* tests were used if the one or both of the previous assumptions were rejected.

## Results

### Larval mortality

Larval mortality was significantly affected by pre-exposure temperature (PET) and test-temperature (TT), with a significant interaction between these two factors. However, larval mortality did not differ significantly between the third and the fifth generation or between replicates. Based on this result, both generations as well as replicates were grouped for further statistical analyses (Table 1).

**Table 1:**
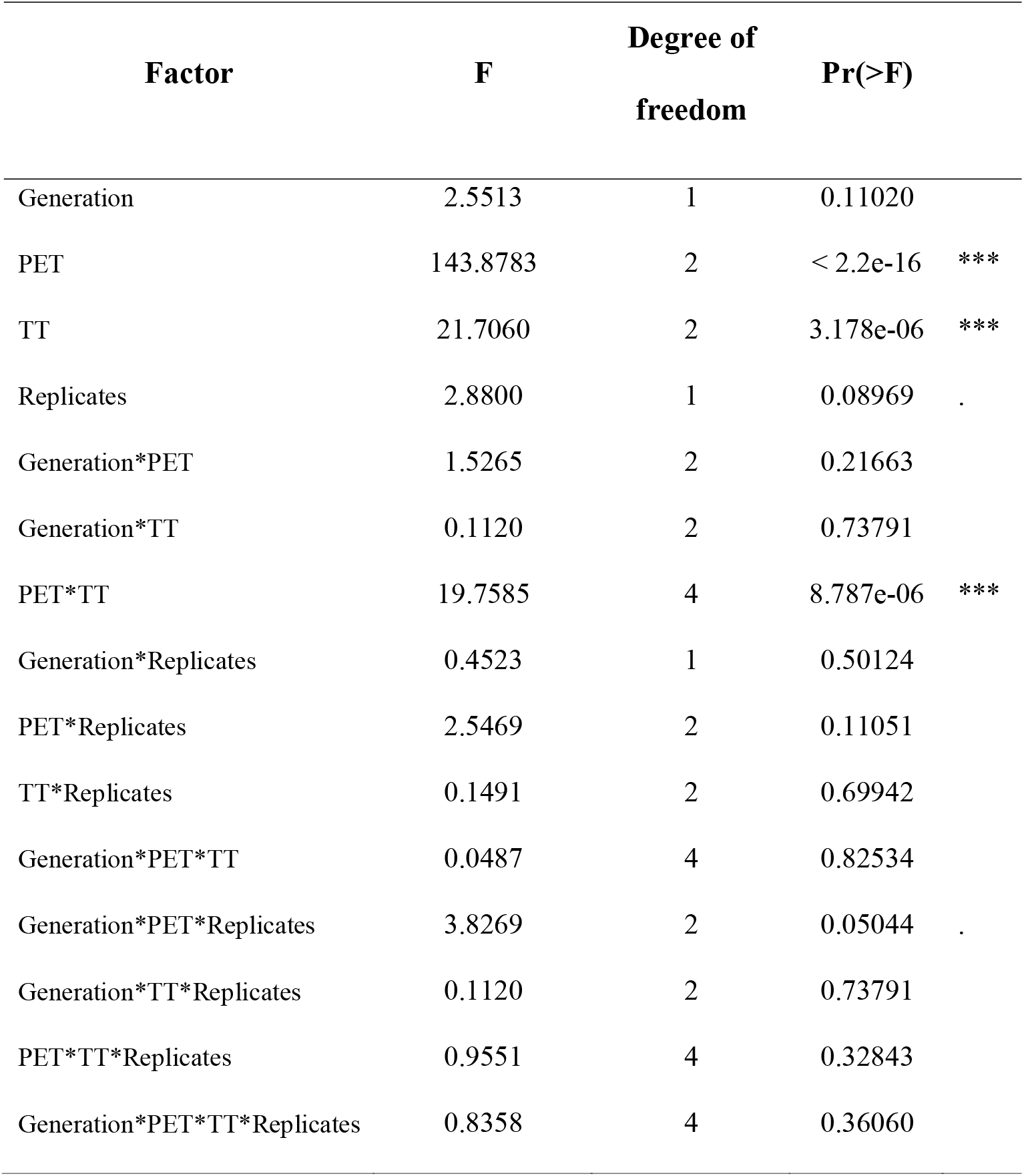
Summary of the linear mixed model executed for the larval mortality after 48h at TT

| Factor | F | Degree of<br>freedom | Pr(>F) |
| --- | --- | --- | --- |
| Generation | 2.5513 | 1 | 0.11020 |
| PET | 143.8783 | 2 | < 2.2e-16 *** |
| TT | 21.7060 | 2 | 3.178e-06 *** |
| Replicates | 2.8800 | 1 | 0.08969 . |
| Generation*PET | 1.5265 | 2 | 0.21663 |
| Generation*TT | 0.1120 | 2 | 0.73791 |
| PET*TT | 19.7585 | 4 | 8.787e-06 *** |
| Generation*Replicates | 0.4523 | 1 | 0.50124 |
| PET*Replicates | 2.5469 | 2 | 0.11051 |
| TT*Replicates | 0.1491 | 2 | 0.69942 |
| Generation*PET*TT | 0.0487 | 4 | 0.82534 |
| Generation*PET*Replicates | 3.8269 | 2 | 0.05044 . |
| Generation*TT*Replicates | 0.1120 | 2 | 0.73791 |
| PET*TT*Replicates | 0.9551 | 4 | 0.32843 |
| Generation*PET*TT*Replicates | 0.8358 | 4 | 0.36060 |

Mean larval mortality increased with the increase in delta temperature according to the temperature off-set between pre-exposure and experimental temperature (ΔT°=TT–PET)(Kruskal-Wallis *χ*^2^_4,175_= 57.253, p-value = 1.1×10-11)(Fig. 2). On the opposite, larval mortality did not decrease concomitantly with a decrease of temperature.

**Fig 2:**
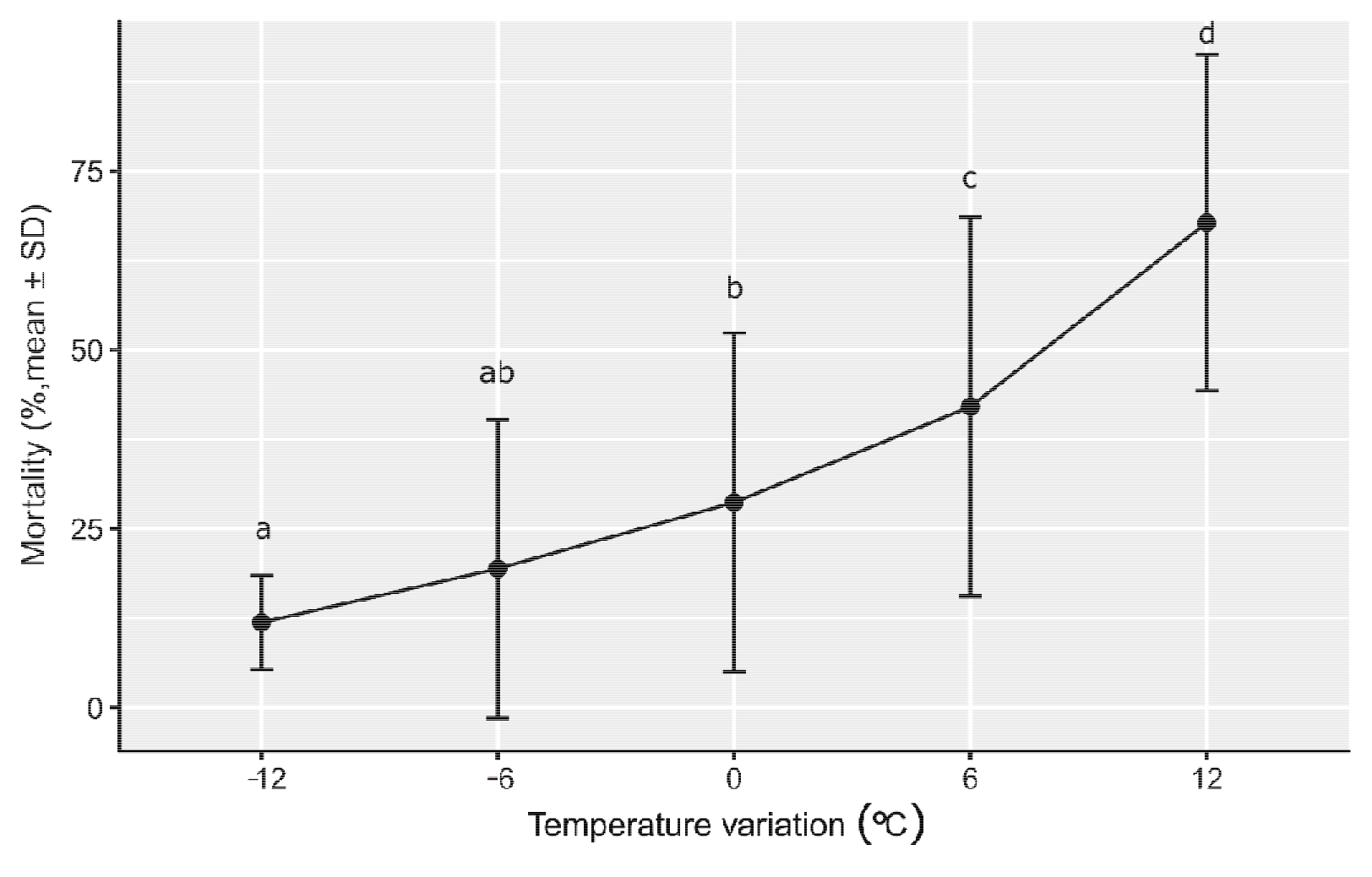
Mean larval mortality after 48h depending on the delta temperature (°C) (delta temperature = TT–PET). Letters denote significant mortality differences between TT (Kruskal-Wallis *χ*^2^_4,175_= 67.320, p-value = 8,347×10^−14^).

After 48 hours, significant differences were found between larval mortality depending on the PET (Kruskal-Wallis *χ*^2^_2,177_= 76,932, p-value <2.2×10^−16^). The overall mortality of the larvae was significantly lower for individuals coming from PET_26°_ compared to a PET_14°_ (*post-hoc* Dunn’s-test p-value = <2×10^−16^) or PET_20°_ (*post-hoc* Dunn’s-test p-value = 7,2×10^−10^) (Fig. 3), independently of TT. By looking at TT depending on the PET, larvae reared at PET_14°_ showed different mortality between TTs (ANOVA F_2,57_= 8,889, p-value = 7,4 ×10^−4^): mortality was significantly higher at TT_26°_ than at both TT_20°_ (*post-hoc* Tukey’s-test p-value = 3,6×10^−5^) and TT_14°_ (*post-hoc* Tuckey’s-test p-value = 54,1×10^−5^) (Fig. 4a). For the larvae from PET_20°_, none of the TT showed a significant difference for larval mortality compared to the PET (ANOVA F_2,57_= 2.936, p-value = 0.06) (Fig. 4b). Finally, for the larvae coming from the PET_26°_, no significant difference was found between the different TT, even though the means showed a trend for lower mortality at higher temperatures (ANOVA F_2,57_= 1.453, p-value = 0.24) (Fig. 4c).

**Fig 3:**
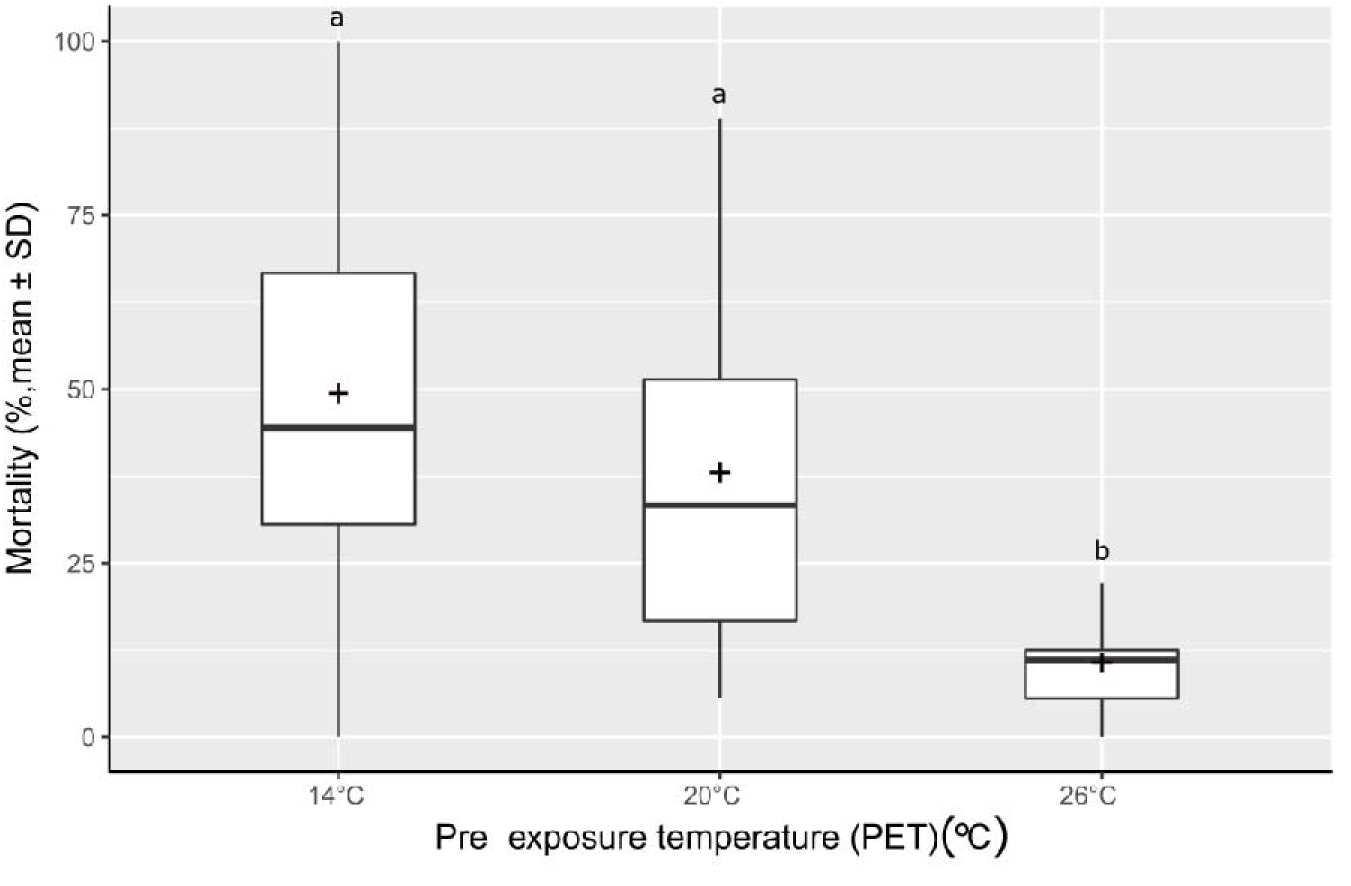
Boxplot of larval mortality after 48h depending on the PET (°C), the crosses show the mean mortality for each PET. The letters denote significant differences between mortality (Kruskal-Wallis *χ*^2^_2,59_= 79.5 1 3, p-value <2.2×10^−16^).

**Fig 4:**
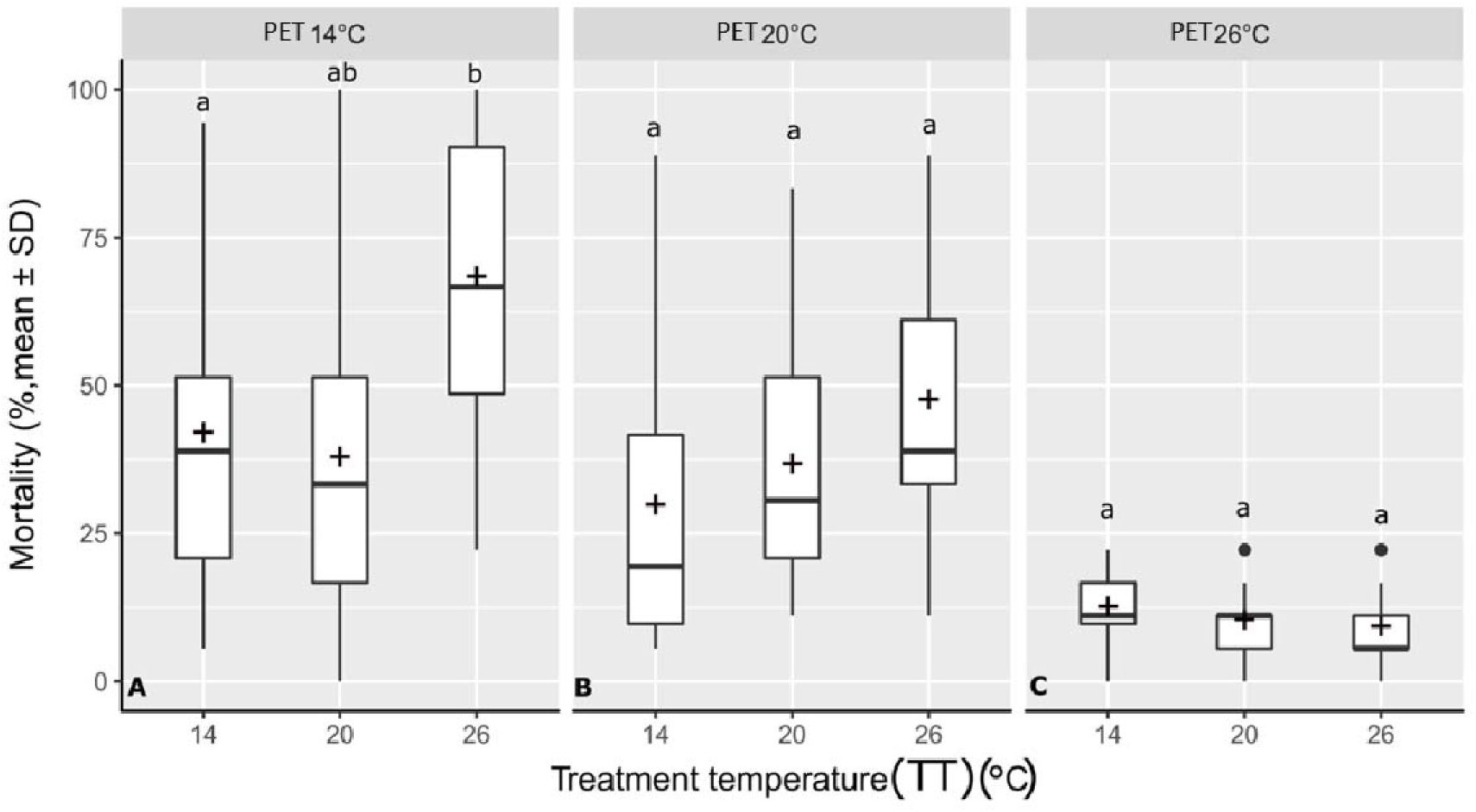
Boxplot displaying larval mortality after 48h by TT (°C) for (A) PET 14 °C (ANOVA F_2,57_= 15,18, p-value = 5,19×10^−6)^, (B) PET 20 °C (ANOVA F_2,19_= 2.936, p-value = 0.06), (C) PET 26 °C (ANOVA F_2,19_= 1.453, p-value = 0.24). Crosses illustrate mean mortality for each PET. Different letters denote significant differences in mortality between groups.

The pupae moratlity was not significantly different between the any of the TTs (ANOVA F_2,177_= 1.023, p-value = 0.36) or PETs (PET_14°_ : Kruskal-Wallis *χ*^2^_2,57_= 2,226, p-value=0,3, PET_20°_ : ANOVA F_2,57_= 0.007, p-value =0,9, PET_26°_: ANOVA F_2,57_= 1.575, p-value = 0.4)(Fig. 5).

**Fig 5:**
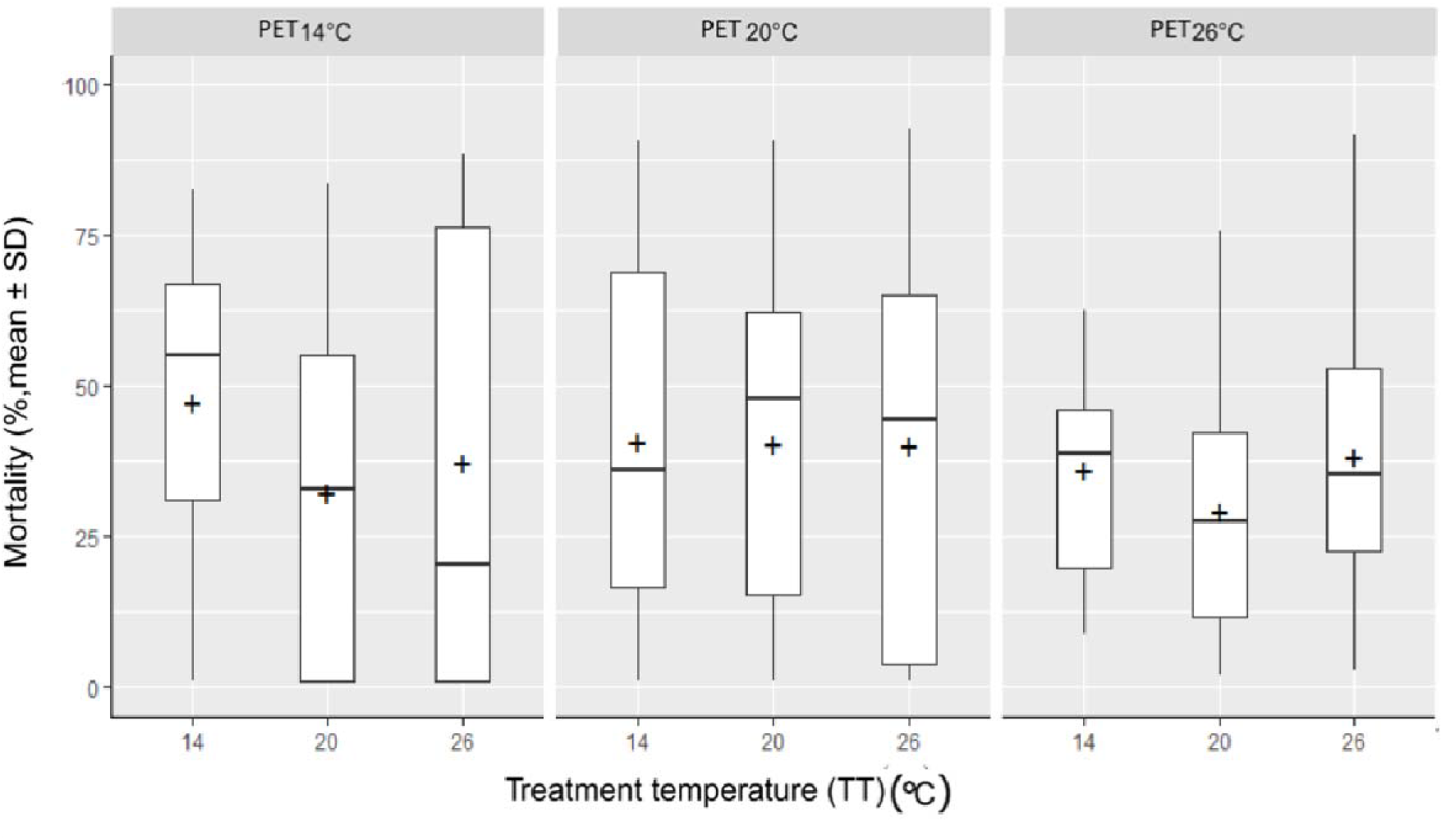
Boxplot displaying the pupae mortality by TT (°C) for (A) PET 14 °C (Kruskal-Wallis *χ*^2^_2,19_= 2,226, p-value=0,3), (B) PET 20 °C (ANOVA F_2,57_= 0.002, p-value =0,9), (C) PET 26 °C (ANOVA F_2,57_= 1.012, p-value = 0.3). Crosses show the mean mortality rate for each PET.

## Discussion

The aim of this study was to investigate the possibility of rapid adaptation of *C. riparius* to seasonal temperature changes. Experimental populations, composed of individuals from five natural populations across a wide distribution range, were reared under three different temperature regimes and mortality rates tested after the third and fifth generation, at the same three temperatures, respectively. The pupation mortality did not display any significant differences between treatments while having an important mortality which is consistent with pupation being the most critical life stage (Oliver, 1971). The insensitivity of pupal mortality to external factors suggests that rather internal factors and processes are responsible for this mortality. For this reason only larval mortality is relevant here. Pre-exposure over three generations to the lowest mean summer temperature experienced by either of the five natural populations the experimental populations were composed of, did not have an influence on larval mortality at low and intermediate experimental temperatures (Fig. 4, left panel) It did, however, increase susceptibility to high temperatures resulting in the highest recorded mortality in our experiments. Apparently, the small number of generations for pre-exposure did not elicit a benign effect on the fitness trait. Negative pleiotropic interactions to heat resistance or transgenerational epigenetic effect (Whittle et al., 2009; Heard & Martienssen, 2014) could have caused the heightened mortality at high temperatures. However, since temperature conditions experienced by the next generation are difficult or even impossible to predict for a multivoltine species, such a canalisation should be selected against if under genetic control. Nevertheless, it is possible that this difference was induced by the experimental protocol, as the experiment did not include feeding during the 48H of treatment. This could have been detrimental for cold raised larvae, because while their developmental rate was increased with high temperature, they lacked the necessary energy. This therefore brings the light on the fact that the higher temperature treatment lead to an increase of development speed. This means that larvae subjected to 26°C treatment theoretically developed more during 48h than at 14°C and this, without food intake

The intermediate temperature regime applied for three and five generations did also not result in a relative or absolute decrease of mortality when the offspring was exposed to this temperature. Probably because 20°C can be considered as the most benign temperature for the species (Supporting information 2), thus presenting no selection pressure large enough to overcome drift in the relatively small experimental populations. The mortality level of 30-40% observed at this temperature is a known base line for the species, at least in experimental conditions (Vogt *et al*., 2007a; Vogt *et al.;* 2007b; Nowak *et al*., 2007a; Nemec *et al*., 2012). The slightly, albeit marginally non-significant increase of mortality at 26°C is potentially due to a stronger selection pressure at this temperature.

The pre-exposure to 26°C yielded surprising results, because it reduced mortality significantly and substantially down to about 12% in all treatment temperatures despite the difference in developmental speed between the treatment temperature (Fig. 4 right panel). The fact that there is no variation among the treatments, i.e. a lack of interaction between the genotype with the respective environment argues for a deterministic, i.e. genetic effect. It is, however, unclear why mortality is not generally so low in *C. riparius*, if it can demonstrably be swiftly achieved by selection. Trade-offs with other unmeasured, costly fitness traits appears a reasonable explanation. Another indication for selective processes was the obvious difference in variance within treatments: in PET_14°_ and PET_20°_, the variance of mortality is high, indicating that a lot of variation is present within replicates, while it is strongly reduced for PET_26°_, which can be expected under selection of resilient phenotypes (Cavicchi *et al*., 1995; Bettencourt *et al*., 2002; Debat & Le Rouzic, 2018). We thus hypothesize that this adaptation is based on standing variation given moderate number of individuals, the low mutation rate of our species (Oppold & Pfenninger, 2017), and the identical response of all replicates.

Our results are consistent with studies in *Drosophila melanogaster*. Populations reared at 28°C survived better when compared to populations reared at lower temperature (Cavicchi *et al*., 1995). The same laboratory populations were later found to have fixed alleles of the heat shock Hsp70 gene, but also a lower level of inducible heat shock proteins at 28°C than at lower TTs (Bettencourt *et al*., 2002). This lower level of inducible heat shock proteins was interpreted as lower capability to respond via phenotypical plasticity to temperature increase (Bettencourt *et al*., 1999); following the hypothesis that in this case Topt has shifted to 28°C. A similar pattern was also found in a natural *D. melanogaster* strain from Africa displaying an exceptional tolerance toward high temperatures (Zatsepina *et al*., 2001). If the adaptation mechanisms are the same in *C. riparius*, they can have an important impact on its population dynamics since climatic models have predicted an increase of temperature by at least 2.6°C. An adaptation to high temperature would then increase their metabolism as well as their developmental speed and thus number of generations without increasing mortality. This could lead to “midge blooms” during summer by increasing population size. This in turn could create problems for human health according to previous findings that Chironomids may be allergenic for humans (Tee *et al.;* 1985, Tee *et al.;* 1987; Eriksson *et al*., 1989) and are able to spread diseases as such as cholera (Broza & Halpern, 2001; Halpern, 2011).

Our results suggest that i) rapid adaptation to different temperature regimes is possible in *C.riparius* within a few generations and ii) therefore the response to seasonal changes in the temperature regime of natural populations may be at least in part be driven by adaptation and not only by phenotypic plasticity. However, our experiments have been conducted on a mixture of populations from across Europe. This mixture perhaps resulted in an artificial increase of genetic diversity composed of warm and cold temperature adapted individuals (Supplementary material 1,2). Therefore, even if it was shown that many of these haplotypes may occur together in natural populations because of high gene flow (Waldvogel *et al*., 2018), the adaptation noticed during our experiment may not be as fast under natural conditions. Still, previous studies indicated that field populations have a high genetic variability (Waldvogel *et al*., 2018). On the other hand, mixing of different populations may have led to outbreeding depression by induction of intrinsic genetic incompatibilities which, in turn, could have slowed or masked adaptation to the external selection regime (Lynch, 1991). Such outbreeding depression has been observed in crossings of different *C. riparius* populations (Oppold *et al*., 2017). In our experiment we only studied mortality as response to temperature; this may have biased our results by overseeing some responses that could have change the overall interpretation of the adaptation mechanisms. Therefore, to strengthen the inference of rapid adaptation over seasonal time scales further, it needs to be shown that similar phenotypic changes also occur in natural populations and that these changes are linked or at least associated with respective genomic allele frequency changes.

## Conclusion

In this study, we showed that *C. riparius* lowered mortality due to high temperature exposure by rapid adaptation rather than phenotypic plasticity. Our results thus indicate that *C. riparius* is able to react to seasonally varying temperature regimes by rapid adaption within a few generations, modifying the survival of the organism not only for warm but also for colder temperatures. This study adds to the currently accumulating literature of rapid adaptation in multivoltine species.

## Acknowledgements

The authors thank D. Lüders for technical assistance. This study was conducted at the Senckenberg Biodiversity and Climate Research Centre Frankfurt.

## Availability of data

The datasets supporting this article are included within the article and its supplementary.

**Compliance with ethical standards**

## Ethics approval

Not applicable.

